# Comparative Analysis of Within-Host Dynamics of Acute Infection and Viral Rebound Dynamics in Postnatally SHIV-Infected ART-Treated Infant Rhesus Macaques

**DOI:** 10.1101/2024.05.21.595130

**Authors:** Ellie Mainou, Stella J Berendam, Veronica Obregon-Perko, Emilie A Uffman, Caroline T Phan, George M Shaw, Katharine J Bar, Mithra R Kumar, Emily J Fray, Janet M Siliciano, Robert F Siliciano, Guido Silvestri, Sallie R Permar, Genevieve G Fouda, Janice McCarthy, Ann Chahroudi, Cliburn Chan, Jessica M Conway

## Abstract

Viral dynamics of acute HIV infection and HIV rebound following suspension of antiretroviral therapy may be qualitatively similar but must differ given, for one, development of adaptive immune responses. Understanding the differences of acute HIV infection and viral rebound dynamics in pediatric populations may provide insights into the mechanisms of viral control with potential implications for vaccine design and the development of effective targeted therapeutics for infants and children. Mathematical models have been a crucial tool to elucidate the complex processes driving viral infections within the host. Traditionally, acute HIV infection has been modeled with a standard model of viral dynamics initially developed to explore viral decay during treatment, while viral rebound has necessitated extensions of that standard model to incorporate explicit immune responses. Previous efforts to fit these models to viral load data have underscored differences between the two infection stages, such as increased viral clearance rate and increased death rate of infected cells during rebound. However, these findings have been predicated on viral load measurements from disparate adult individuals. In this study, we aim to bridge this gap, in infants, by comparing the dynamics of acute infection and viral rebound within the same individuals by leveraging an infant nonhuman primate Simian/Human Immunodeficiency Virus (SHIV) infection model. Ten infant Rhesus macaques (RMs) orally challenged with SHIV.C.CH505 375H dCT and given ART at 8 weeks post-infection. These infants were then monitored for up to 60 months post-infection with serial viral load and immune measurements. We use the HIV standard viral dynamics model fitted to viral load measurements in a nonlinear mixed effects framework. We find that the primary difference between acute infection and rebound is the increased death rate of infected cells during rebound. We use these findings to generate hypotheses on the effects of adaptive immune responses. We leverage these findings to formulate hypotheses to elucidate the observed results and provide arguments to support the notion that delayed viral rebound is characterized by a stronger CD8+ T cell response.

## 1 Introduction

HIV-1 continues to pose a significant global health challenge, affecting 38.4 million individuals as of 2021, according to the World Health Organization [1]. Among these, 1.7 million are children, as reported by UNAIDS [2]. Most of these children live in sub-Saharan Africa and were infected perinatally through breastfeeding [3].

The course of HIV infection and the response to treatment in infants and children differ significantly from those observed in adults. Infants experience a rapid increase in plasma viremia, reaching levels much higher than adults [4, 5]. The decline in infant plasma viremia is slow, reaching a setpoint at around 5 years of age [6, 7]. Furthermore, the rates of clinical disease progression vary between adults and children. For instance, 20-30% of children undergo rapid progression to AIDS or death in the first year of life [6]. These rates of disease progression also vary based on the age of the child; a 1-year-old with CD4+ T cells being 10% of total lymphocytes has a 40% risk of progressing to AIDS within a year, while a 10-year-old with the same laboratory values has a 7.4% risk within the same time frame [6, 8]. Pediatric elite controllers are also rarer; only 0.008% of HIV-infected children are elite controller, in comparison to 0.5% of adults [3, 9, 10]. These distinctions are primarily attributed to differences in the immunologic development of infants and the rapid expansion of CD4+ T cells associated with somatic growth [6].

Immune responses are age-dependent [6,11]. This dynamic environment of the developing immune system is believed to contribute to the differences in clinical outcomes both between infants and adults and among infants infected at different stages of their development [6]. Adaptive immune responses undergo development with age, commencing around 7–9 weeks of gestation when T-cell progenitor cells colonize the thymus [12, 13]. B lymphocytes are also detectable by the end of the first trimester of gestation [6, 14]. Neonatal T cells are predominantly naive, with reduced proliferative responses and lower IL-2, and IFN*α*-IFN*γ* secretion. CD4+ T cells in neonates have diminished CD40L expression and provide less support for B-cell function [6, 15]. Circulating CD4+ T cells are significantly higher in infants and children, peaking at 3–4 times adult levels in the first months and gradually declining to adult levels by age 6 [16]. Variability in CD8+ T cell numbers is less pronounced in children and infants compared to adults [17]. Regarding humoral immunity, naive infant B cells express lower levels of CD21, CD40, CD80, and CD86. Neutralizing antibodies develop at least as commonly as in adults and in some cases can be detected within one year of infection [18]. In addition, neonatal antibody responses have lower peak levels, and persist for a shorter duration [6, 15]. These antibodies exhibit less affinity maturation, heterogeneity, and differ in IgG isotype distribution compared to adults [6, 15]. Immunoglobulin levels, especially IgA, remain lower than in adults, even at one year of age [3, 19]. However, broadly neutralizing antibodies, whose development is the goal of current vaccine approaches, arise earlier in pediatric infections compared to adults (1 year compared to 2-3 years) and show higher breadth and potency [18, 20–24]. Overall, in utero and early life, immune development establishes a highly tolerogenic and broadly anti-inflammatory environment [3], influencing the response to HIV infection.

Studies on pediatric immunity to HIV indicate insufficient responses to control the infection. For instance, HIV-specific CD4+ T-cell responses are infrequent in perinatally infected infants under 3 months of age and remain low even with early ART initiation [6, 25, 26]. HIV-specific CD8+ T-cell responses can be detected at birth for in utero infections [27]. Acute infection in infants is characterized by a response to env, whereas in chronically-infected infants CD8+ T cell responses targeted mainly gag, pol and nef proteins [27]. Additional studies demonstrate that CD8+ T cells are vital for HIV control in children too: pediatric elite controllers (ECs), compared to progressor counterparts, have a higher frequency of HIV Gag-specific, polyfunctional CD8+ T cells capable of cytokine release [3, 10, 28]. Regarding humoral responses, active production of HIV-specific ADCC antibodies is observed in most of HIV-infected infants only after 12 months of age [6, 29]. There is also evidence for the potential role of non-neutralizing HIV-specific antibodies in protecting against disease progression and viral control. Muenchhoff et al. noted that a subset of ART-naïve pediatric non-progressors had higher p24-specific IgG levels than children under ART [3, 30]. Overall, these studies suggest that, despite a developing immune system, infants can mount an immune response against HIV, but that response is not effective at controlling the infection, leading to continued replication of the virus [27].

Innate immunity is a crucial part of the infant immune system since, given the limited exposures to antigens in utero, infants must rely on their innate responses for their protection [31–34]. Innate immune cells in early life exhibit distinctive cytokine profiles, characterized by lower levels of proinflammatory cytokines and higher levels of immunoregulatory and anti-inflammatory cytokines, fostering a tolerogenic environment [35]. Neonatal NK cell frequencies are comparable to those in adults [13,36]. However, NK cells display differences in the expression patterns of inhibitory and activating cell surface receptors [13,37]. NK cells from vertically HIV-infected children exhibit decreased cytolytic activity compared to uninfected children and reduced ADCC capacity compared to adults [13, 38–41]. Though generally viewed as nonspecific innate effector cells, mounting evidence suggests that NK cells possess adaptive, memory-like characteristics [42]. In humans, a subset of these long-lived NK cells with memory-like traits has demonstrated heightened effector responses when encountering viral antigens again or when properly activated by pro-inflammatory cytokines [43–48]. Overall, features of innate immunity are shared with adaptive and thus innate immunity may be harnessed for enhanced protection [49].

Mathematical modeling has played a crucial role in understanding HIV post-treatment control and viral rebound dynamics after treatment interruption [50–56]. Various studies have employed mathematical models to estimate recrudescence rates and investigate inter-individual heterogeneity in viral rebound dynamics [51–53]. Mathematical modeling has also been used to explore the effect of latent reservoir size on rebound times and the reduction needed in the reservoir [50] or the strength of immune responses to achieve a cure in adult humans [57, 58]. However, there have not been modeling efforts to compare the dynamics of acute infection and post-treatment interruption, especially in the same pediatric individuals.

In this study we aim to determine the differences in the within-host HIV viral dynamics between acute infection and after treatment interruption in the same pediatric individual. Understanding the initial stages of HIV infection in children, as well as the progression of the infection within the host may be crucial for the identification of factors for the development of an effective vaccine or for viral control in children. However, performing studies on human infants is difficult. Non-human primate models of Simian Immunodeficiency Virus (SIV) and Simian/Human Immunodefiency Virus (SHIV) display many similarities to human infections and have provided valuable insight [6, 59–62]. Here, we use experimental data from an infant nonhuman primate Simian/Human Immunodeficiency Virus (SHIV) infection model that mimics breast milk HIV transmission in human infants. Infant rhesus macaques (RMs) were orally challenged with SHIV.C.CH505 375H dCT 4 weeks after birth and ART was initiated at 8 weeks post-infection (wpi) and terminated a year later. We employ the standard HIV viral dynamics model [63, 64], which is a simple and parsimonious model that accurately describes within-host dynamics. We focus on a simple model, as we are interested in describing the broad differences between acute infection and viral rebound. We fit the standard model to viral load measurements of acute infection and post-analytic treatment interruption (ATI) of 10 RMs in a nonlinear mixed effects (NLME) framework, using the stage of the infection as a covariate (i.e. assume different values of a parameter based on the stage of the infection) across all combinations of parameters, to identify differences between acute infection and rebound dynamics. Our goal is to capture the broad differences between acute infection and viral rebound through the values of the standard model’s parameters and then generate hypoteheses that could explain those difference. We also do not select a single model that best fits our data but focus on an ensemble of equivalent models and their common features. We find that the main difference between acute infection and viral rebound lies in parameters that are related to cellular immune responses, both cytolytic and non-cytolytic.

## 2 Methods

### 2.1 SHIV infection data in infant macaques

Ten infant *Rhesus macaques* (RMs) were orally challenged with SHIV.C.CH505 four weeks after birth. Viral load measurements were taken weekly. Antiretroviral treatment (ART) was initiated at 8 weeks post-infection (wpi) and was maintained for one year. After treatment interruption, subjects were followed for viral rebound, with regular viral load measurement taken every 2-3 days (Fig 1). The viral detection threshold is 60 viral copies/mL. The experiment is described in detail in [65, 66]. In addition to viral load, potency of antibodies in neutralizing autologous virus envelope glycoprotein gp120 was measured by a TZM-bl assay [67] was measured on the day of treatment interruption.

**Figure 1:**
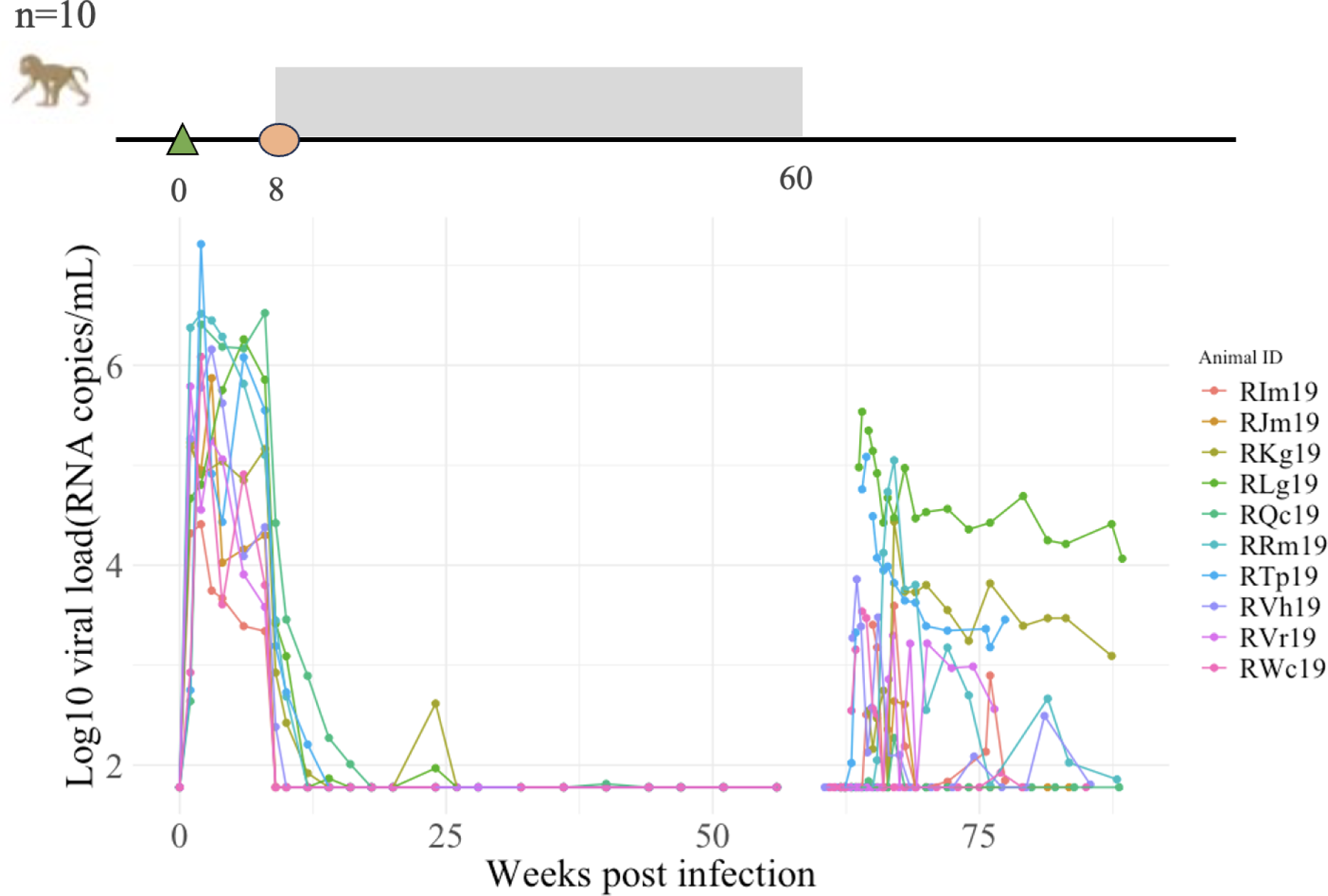
Description of the experiment. Study schematic showing ART timing (grey shaded area), as well as viral load trajectories for the 10 subjects in the experiment.

### 2.2 The model

We employ the standard viral dynamics model, initially devised to investigate viral decay following the initiation of treatment [64]. This model, known for its ability to encapsulate within-host dynamics of HIV infection, has been deliberately chosen for its simplicity and parsimony. The emphasis on a straightforward model serves as an approach to delineate the broad differences in dynamics between acute infection and viral rebound. The standard model is a target cell-limited model, i.e. HIV infection is limited by the availability of target cells, consisting of three compartments — uninfected cells, infected cells, and free virus. Target cells, *T*, are produced at rate *λ*, die at rate *d* and become infected at rate *β*. Infected cells, *I*, die at a constant rate *δ* and produce HIV-1 virions at rate *p*. Free virus, *V*, is cleared from the blood at rate *c* (Fig 2). This model can be expressed as the following system of differential equations:

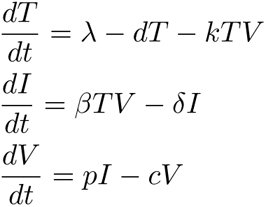

**Figure 2:**
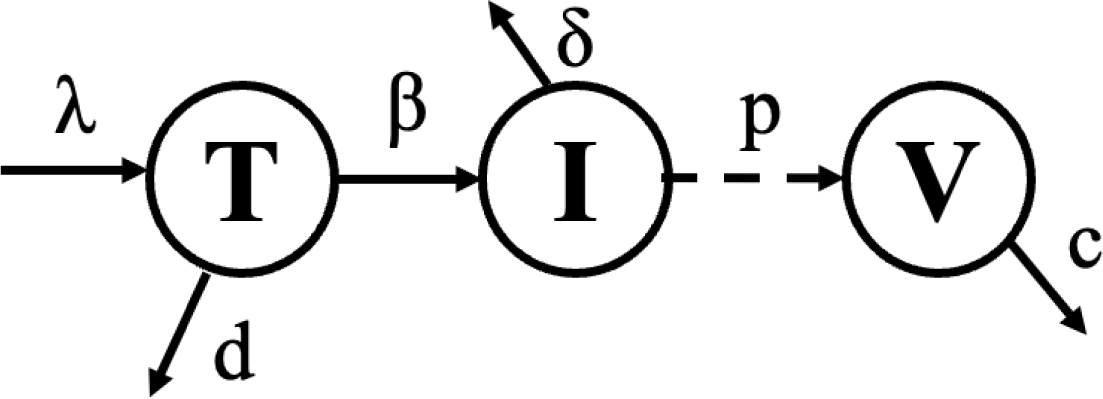
Standard viral dynamics model schematic. A) Schematic of the standard model. Target cells, *T* are sourced at rate *s* and die at rate *d*. Virus *V* infects target cells at rate *β*. Infected cells, *I* die at rate *δ* and produce virions at rate *p*. Free virus gets cleared at rate *c*.

This target-cell limited model has been shown to capture the dynamics of early infection [68]. We focus on a simple model to characterize the general differences between the different groups. Because we are interested in broad differences between the two groups, the standard model, without incorporating explicit dynamics of adaptive immune responses, is still appropriate for capturing the dynamics of viral rebound. Adding explicitly immune responses in the model, would increase the number of parameters, thus adding uncertainty. The standard model exhibits both simplicity and parsimony [69].

We rescale *T* and *I* by *T*_0_, where *T*_0_ is the initial number of target cells per mL. This restructuring makes plain that the viral production rate *p* and *T*_0_ structurally unidentifiable and permits us to estimate the single composite parameter *pT*_0_. We assume that prior to infection, target cells are at equilibrium and therefore, we set *λ* = *d · T*_0_ [68]. We also fix *d* = 0.01 per day [68]. We estimate five parameters– *β*, *δ*, *p*, *c*, as well as *t_start_*, the onset of exponential viral growth, by fitting the model to plasma viral load data during acute and rebound infection.

### 2.3 Model Fitting

We fit the standard viral dynamics to viral load measurements during acute infection before treatment and after treatment interruption. We are particularly interested in elucidating the differences between the two groups and obtain a more parsimonious model. For this reason, we use a nonlinear mixed effects (NLME) framework, as this approach allows us to capture both population-level effects, as well as individual variability. We test all models in which the stage of the infection (acute or rebound) acts as a covariate affecting all possible combinations of parameters (a total of 31 combinations). This means that for the parameter for which the stage of infection is a covariate, we assume that the values of that parameter for acute infection come from distributions of different means. We fix the population-level value of the viral clearance rate to c=3, 6 and 12 d*ay^−^*^1^ [64, 68, 70], while still allowing for inter-individual variability. To better explore the parameter space, we test each model using the best initial parameter value guess along with four additional sets of initial parameter values selected randomly from an interval around the original value. We also add correlations between random effects in pairs of parameters when ANOVA tests indicate significance at the 0.01 significance level. Therefore, with 31 possible covariate combinations, 3 fixed values for the viral clearance rate and 5 initial sets of parameter values, we end with 465 models to test. In addition, we test 154 models with added correlations between random effects, resulting in a total of 619 models (Fig 3). We run all models in *Monolix* through the *lixoftConnectors* package in *R*.

**Figure 3:**
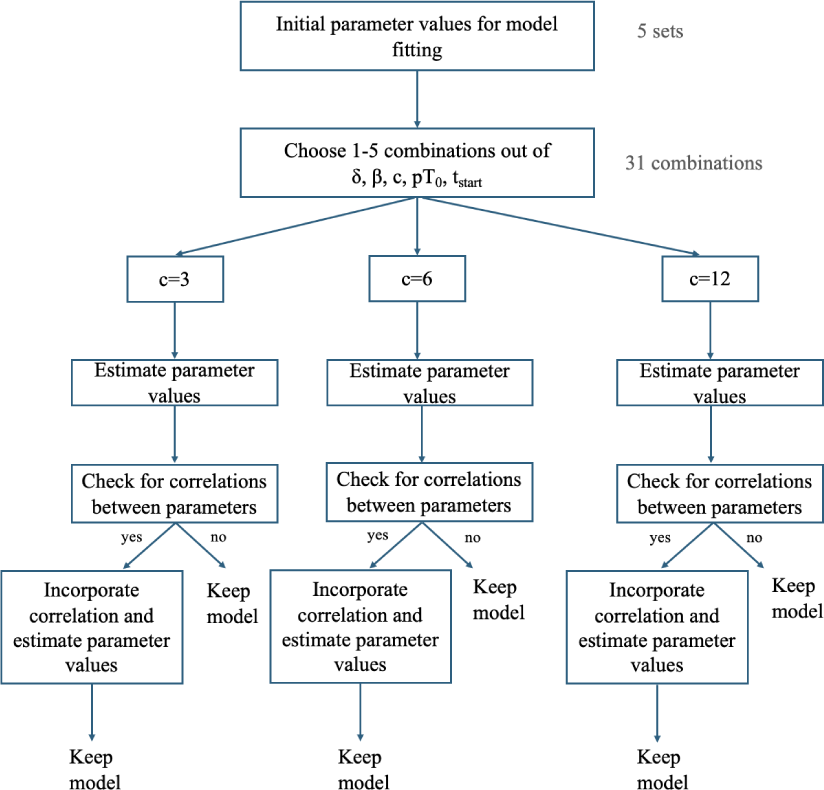
Model Fitting Schematic. Schematic of the approach we follow for model fitting and types of models tested.

### 2.4 Model Selection

We do not select a single model that best fits our data. Instead, we focus on an ensemble of models that are comparably successful in explaining the observed data. Determining the common features of the ensemble provides robustness to the observed features. In addition, we do not focus on a single metric for model selection, namely the Akaike Information Criterion (AIC), which is the standard method for model selection. AIC primarily takes account of the sampling errors in parameter estimation and can offset errors in the estimation of model parameters; the more parameters a model has, the more these methods can compensate, thus balancing model fit with model simplicity, where simplicity is defined by fewer parameters. However, when choosing a model, it’s important to consider its ability to generalize predictions to different contexts. Therefore, we are interested in a more comprehensive approach, that includes the model’s predictive ability, as well as the stability of parameter estimates. We determine these models using the following criteria:

1. AIC: we consider models whose AIC is within 10 points from the lowest AIC.
2. Visual Predictive Check (VPC): The VPC is a graphical method used in statistical modeling to assess the goodness-of-fit and predictive performance of a model. It is a valuable tool for evaluating how well a model captures the observed data and how well it can predict future observations. We focus on models where the summary statistics from the observed data fall within the range of values obtained from the simulated datasets, which indicates that the model can replicate the observed data well.
3. Stability of parameter estimates: We evaluate the % rise in standard error (RSE), which is a statistical measure that quantifies the increase in the standard error of a parameter estimate when comparing two different models or conditions. The RSE is a useful metric that helps assess whether changes in the model’s complexity or data have a substantial impact on the reliability of parameter estimates. To assess the quality of a model RSE, we devise a metric titled *S* defined as the sum of RSE values above 50% divided by the number of parameters estimated.

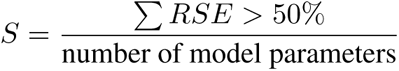

For example, for the model estimates in Table 2, the estimated *S* value would be

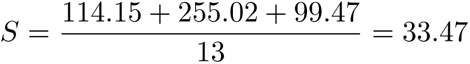

We first focus on models with AIC relative to the lowest, Δ*AIC <* 10. No additional models were excluded upon examination of the visual predictive check, as they displayed no areas of model misspecification. To narrow down our selection, we further restrict Δ*AIC* values, and we choose models with *S <* 60, and Δ*AIC <* 8 or Δ*AIC <* 4. It should be noted that allowing for an increased *S* threshold (*S <* 80) introduces inconsistency in the predicted association between the stage of the infection and the mean parameter values with one model predicting a decreased population value of the viral clearance rate and the onset of viral exponential growth (Supplementary Fig 1). This observation offers support for keeping *S* threshold low.

## 3 Results

We fit the standard viral dynamics model to viral load measurement taken before treatment initiation and after treatment interruption in a nonlinear mixed effects (NLME) framework. The resulting model fits yield varying AIC values. We first focus on models with AIC relative to the lowest, Δ*AIC <* 10, which results in 59 models (Fig 4). No additional models were excluded upon examination of the visual predictive check, as they displayed no areas of model misspecification. To narrow down our selection, we further restrict Δ*AIC* values, and we choose models with *S <* 60, and Δ*AIC <* 8 or Δ*AIC <* 4 resulting in a collection of 13 and 5 models respectively. The results are qualitatively equivalent and below we report the specific results for Δ*AIC <* 8, since it is larger set.

**Figure 4:**
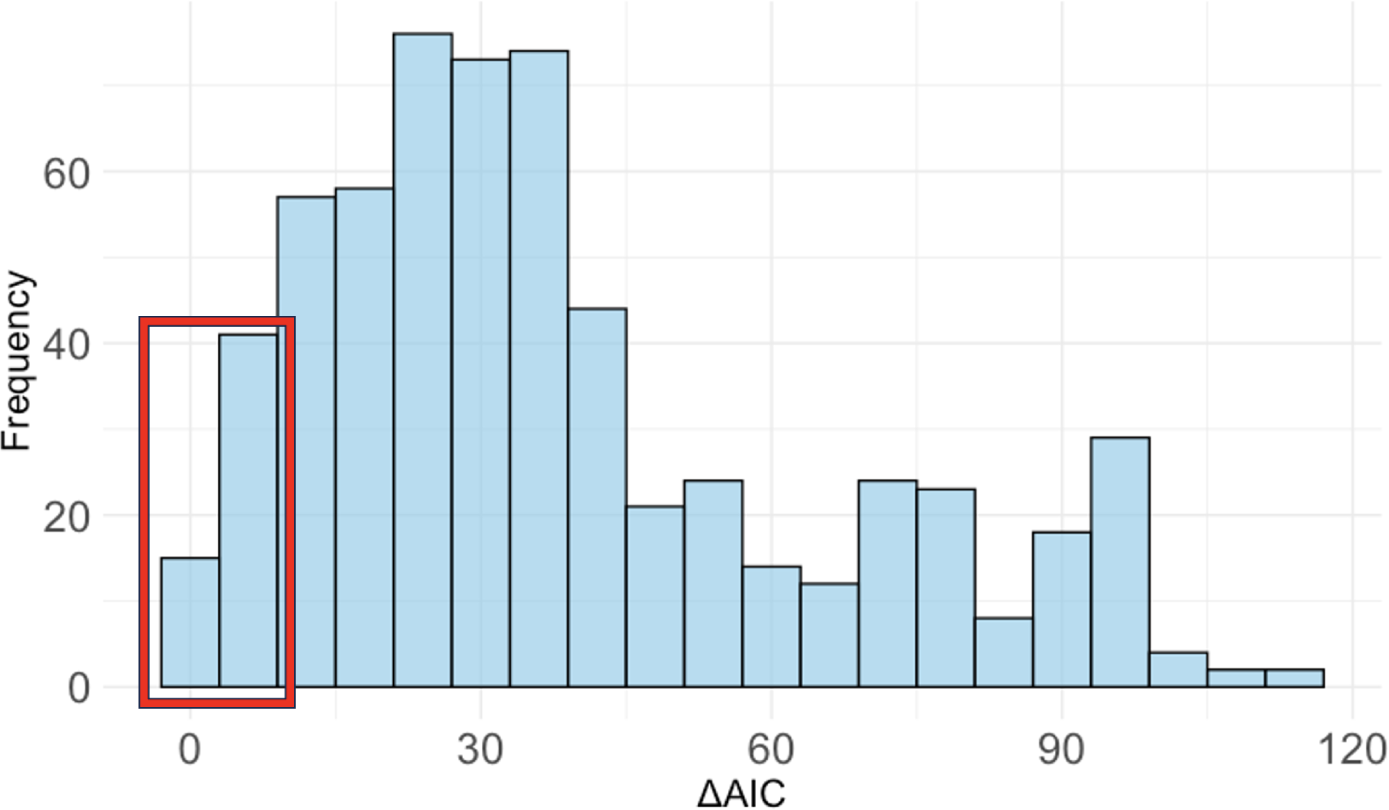
Distribution of Δ*AIC*. Histogram of AIC values relative to the lowest AIC of all the models tested. We reject models with Δ*AIC >* 10.

Table 1 provides a summary of the features of the 13 selected models. Below, we report the common features among selected models; for example, how many of those 13 models predict a difference in the mean value of a specific parameter between acute infection and viral rebound and does the parameter value increase or decrease for rebound dynamics (Fig 5)? We focus on the population-level averages of our model parameters, where the two populations refer to acute infection and viral rebound. We will compare the differences in parameter values between acute infection and viral rebound and generate hypotheses for potential mechanisms that these differences can be attributed to.

**Figure 5:**
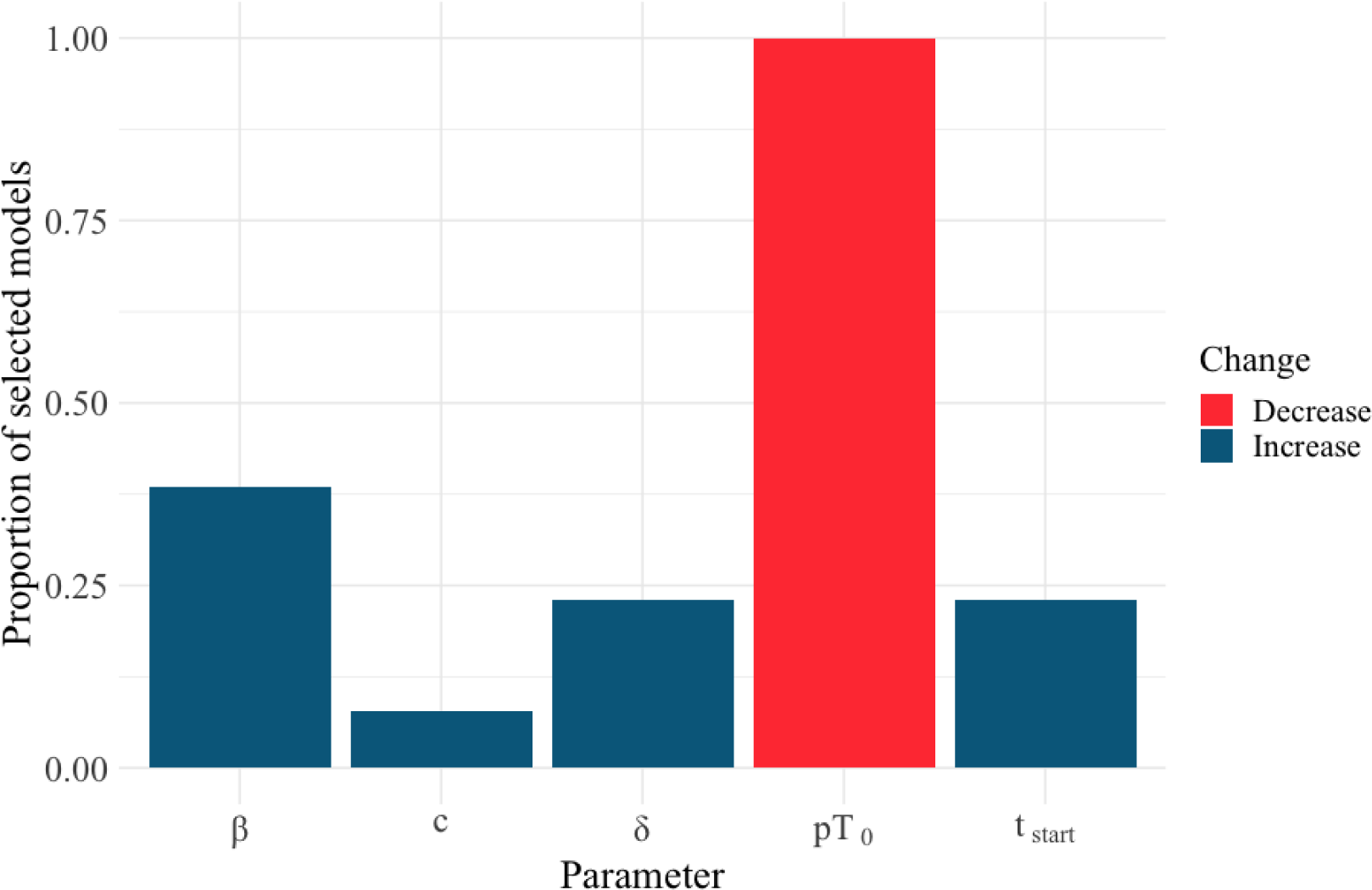
Parameters differing between acute infection and viral rebound. Bar chart depicting the proportion times of a parameter is predicted to differ between acute infection and viral rebound in all selected models. Viral clearance rate (*c*) is predicted to be different between acute infection and viral rebound in 1 out of 13 best models; death rate of infected cells (*δ*) and start of exponential viral growth (*t_start_*) in 3 out 13 models; mass-action infectivity (*β*) in 5 out of 13 models and the product of viral production rate (*p*) and start of initial number of target cells (*T*_0_) in all 13 models. Red slices signify a decrease in the value of the parameter for rebound, whereas blue-toned slices depict an increase in the rebound value of the parameter.

**Table 1:**
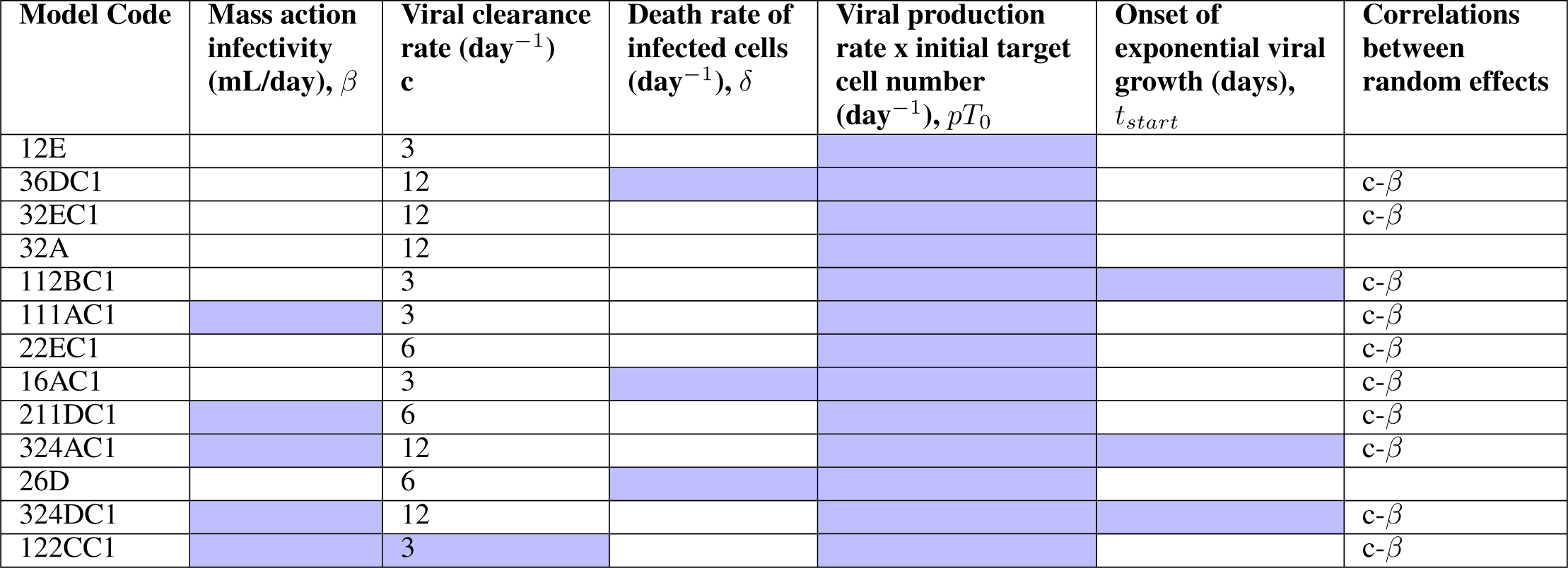
Summary of selected models. For each of the selected models, the blue-colored cells signify the parameters for which we assume a difference in the mean value between acute infection and viral rebound. We also report the mean viral clearance rate *c*, as well as the for which parameters we incorporate correlations between random effects.

**Table 2:** Parameter values: estimates and rise in standard error (RSE) for one of the models selected (also Supplementary Fig 2). In this model we fix the median viral clearance rate to c=3 d*ay^−^*^1^for acute infection, and we assume that the product of viral production rate and initial number of target cells (*pT*_0_) as well as the mass action infectivity (*β*) are difference between acute and rebound and add a correlation between *c* and *β*. RSE values between 50-100% are shown in blue, values between 100-200% are shown in orange an above 200% in red.

|  | Value | RSE % |
| --- | --- | --- |
| Fixed effects |  |  |
| $\delta$ | 0.17 | 32.78 |
| $\log_{10}(pT_0)$ | 6.41 | 3.57 |
| $b_{\log_{10}(pT_0)}(\text{Rebound})$ | -2.1 | 22.77 |
| $c$ | 3 | |
| $\log_{10}(\beta)$ | -4.94 | 11.17 |
| $b_{\log_{10}(\beta)}(\text{Rebound})$ | 1.18 | -16.92 |
| $t_{start}$ | 3.11 | 114.15 |
| Standard deviation of the random effects |  |  |
| $\delta$ | 0.86 | 12.07 |
| $\log_{10}(pT_0)$ | 0.21 | 14.11 |
| $c$ | 1.9 | 255.02 |
| $\log_{10}(\beta)$ | 0.47 | 27.77 |
| $t_{start}$ | 0.8 | 99.47 |
| Correlations |  |  |
| $c - \log_{10}(\beta)$ | 0.92 | 11.27 |
| Error model parameters |  |  |
| $a$ | 0.58 | 4.57 |

### Exponential viral growth phase is initiated later in viral rebound

Out of the 13 models selected, three of them predict that the mean value for the onset of exponential viral growth (*t_start_*) is slower during rebound compared to acute infection (Fig 5). This observation is expected, since the development of immune responses during rebound can reduce viremia, thus delaying rebound [71]. However, for the remaining 10 selected models, such a difference in mean values is not predicted. Instead, distribution of *t_start_* values for acute infection and viral rebound tend to overlap (Supplementary Fig 3), pointing to the inherent heterogeneity in the stochastic phase between treatment interruption and onset of exponential viral growth.

### Humoral immunity appears underdeveloped

Counter-intuitively, only one out of 13 models predicts a difference in mean viral clearance rate between viral rebound and acute infection (Fig5). Since the effects of adaptive immune responses are thought to be more pronounced during rebound dynamics, we would expect neutralizing antibodies to increase the viral clearance rate [72]. Our modeling results do not provide a strong signal for that effect. However, the distribution of viral clearance values for rebound is wider compared to acute infection and allows for values consistent with adult rebound values [57, 73]. Therefore, even though the selected models predict no difference in the mean viral clearance rate, there is high heterogeneity in clearance rate values rebound values (Supplementary Fig 4). The adaptive immune response mechanism that could explain increased viral clearance rate is neutralizing antibodies (nAbs). However, we believe that this heterogeneity cannot be attributed to nAbs, since we do not observe a difference in the rebound viral clearance rate between the RMs that developed neutralizing responses and those that did not (Supplementary Fig 5). Instead, we hypothesize that the heterogeneity in the viral clearance rate may be explained by viral evolution [74]. Furthermore, 5 out of 13 models predict an increase in the viral infectivity rate by approximately one order of magnitude (Fig 6B).

**Figure 6:**
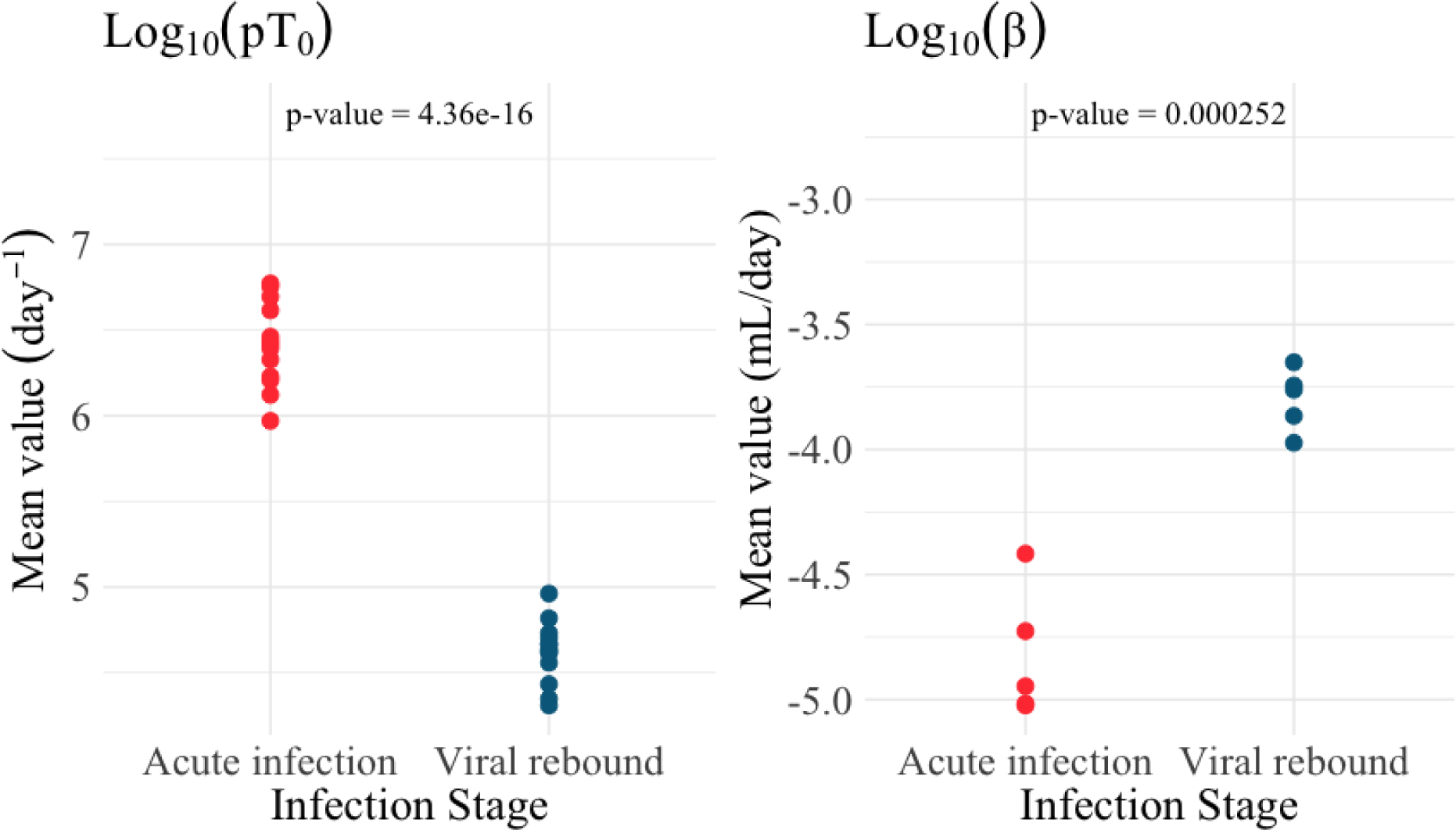
Difference in mean parameter values between viral rebound and acute infection. Mean value for the product of viral production rate and initial number of target cells, *pT*_0_ (A) and mass action infectivity *β* (B) for acute infection (red) and viral rebound (blue) for the 13 models selected. For both parameters, the difference is mean values is statistically significant (T test, p<0.001).

### Viral rebound is characterized by increased killing rate of infected cells albeit with heterogeneity

All the selected models predict a decrease in the mean value of the product of the viral production rate and the initial number of target cells, which are structurally unidentifiable and thus estimated as one parameter. Furthermore, for a subset of selected models (3 out of 13) the mean death rate of infected cells is higher during viral rebound compared to acute infection (Fig 5). In all selected models, the distribution of the estimated death rate of infected cells is wider than for acute infection (Supplementary Fig 6), suggesting that even in these models, there are some RMs that develop these cytotoxic responses. There are multiple mechanisms that could explained an increased infected cells death rate. These include the effect of cytotoxic T lymphocytes (CTLs), NK cells, antibody-dependent cell-mediated cytotoxicity (ADCC) and antibody-dependent cell-mediated phagocytosis (ADCP). We believe that a likely explanation is CTL responses; RMs that developed neutralizing responses also developed ADCC responses (data not shown), but we do not observe a difference in estimated death rate of infected cells (Supplementary Fig 7, Wilcoxon signed rank test p-value *>* 0.05 for all selected models).

## 4 Discussion

In this study, we use the standard viral dynamics model fitted in a NLME framework, to investigate the differences between acute infection and viral rebound in the same SHIV-infected infant RMs. We do not focus on viral rebound times, but on the downstream dynamics after rebound. We are also interested in determining the broad differences between these two stages of the infection, as expressed by the parameters of the standard model and then generate hypotheses where these differences can be attributed to. We use the standard viral dynamics model to provide a broad view and generate hypotheses on differences rather than focusing on specific mechanisms. We find that the main difference in the dynamics is the increased death rate of infected cells and decreased viral production rates and availability of target cells, pointing to the effect of cytotoxic T lymphocytes. Overall, our modeling results suggest changes in parameter values that are associated with a more pronounced cellular response compared to humoral responses.

It should be noted that the differences we find between acute infection and viral rebound are associations with parameter values. Even though this provides a simple framework to explore patterns in the relationship between the stage of the infection and key viral kinetics parameters, the mechanisms that account for those relationships may sometimes be hard to parse out. For example, we find increased death rate of infected cells, which we attribute to CD8+ T cell responses. There is a growing body of evidence that suggests that CD4+ T cells also have a direct killing effect [75, 76]. Similarly, natural killers (NK) cells also have cytotoxic effects and have been shown to have an antiviral effect in HIV infection [77]. An important future direction would include developing models that incorporate more explicit mechanisms, along with longitudinal measurement of the compartments of interest to further elucidate these dynamics.

To draw our conclusions regarding differences between acute infection and viral rebound dynamics, we adopt a novel strategy for model selection. Instead of relying solely on the Akaike Information Criterion (AIC), which is considered the standard for choosing the most parsimonious model, we opt for a diverse set of metrics and diagnostics to identify a set of optimal models. This approach allows for a more comprehensive assessment of model fits that includes model misspecification and stability of parameter estimates. Alongside AIC, we introduce a metric called *S* to evaluate the stability of parameter estimates and utilize the Visual Predictive Check (VPC), which assesses misspecifications in structural, variability, and covariate models. Our model selection involves considering various threshold combinations for each metric (ΔAIC<8 vs ΔAIC<4 and *S*<60 vs *S*<80). We observe that the Δ*AIC* thresholds used in the analysis yield consistent effects for the change in parameter values. However, increasing the *S* threshold from 60 to 80 introduces some inconsistency: we have one selected model predicting a lower viral clearance rate for rebound and another selected model predicting a faster onset of exponential viral growth. These observations offer support for maintaining a lower threshold for our *S* metric.

We find that even though the majority of selected models do not predict a difference in the mean viral clearance rate, the distribution for rebound is wider, allowing for values consistent with adult human rebound [57]. Furthermore, 5 out of 13 models predict an increase in the viral infectivity rate by approximately one order of magnitude (Fig 6B).This observation contrasts other experimental data [78] and mathematical modeling results [79]. We speculate that the increased viral infectivity rate during rebound could be attributed to the increased proportion of target cells that can be easily infected. HIV-specific activated CD4+ T cells are the preferential target of infection, and undergo a clonal expansion due to stimulation by the virus [80–82]. This means that at later stages, there is a larger proportion of target cells that can be easily infected, thus increasing the viral infectivity rate. In addition, all of the selected models predict a decrease in the mean value of the product of the viral production rate and the initial number of target cells. Since *p* and *T*_0_ are not identifiable, we cannot determine which parameter accounts for this decrease. We hypothesize that either parameter decreases. Regarding *T*_0_, in pediatric HIV, CD4+ T cells do not recover to pre-infection levels under ART [3, 83, 84], which could lead to a decreased rebound value. Viral production rate could also be lower at later stages of the infection due to non-cytolytic effects of CD8+ T cells.

Our results point to the importance of CD8+ T cell responses [27, 85] and indicate the development of both cytolytic and non-cytolytic effects during rebound. We observe an increase in the killing rate of infected cells, although the estimated death rate is generally lower compared to the adult rebound values [57, 86] and the variability among rebounding RMs, some of whom display values consistent with acute infection (Supplementary Fig 5). In addition, we find a decrease in the product of scaled viral production rate, which can be attributed to a reduced viral production due to noncytolytic effects of CD8+ T cells [87]. Recent modeling works also provides support for a dual function of CD8+ T cells in HIV-1 infection [87]. Indeed, CD8+ T cell responses are a hallmark of control. For instance, in pediatric, as well as adult, elite controllers display a higher frequency of HIV Gag-specifc, polyfunctional CD8+ T cell responses capable to cytokine release compared to progressors [3, 88, 89]. Experimental findings also support this observations: CD8+ T cell depletion in nonhuman primate models of HIV/SIV/SHIV resulted in a more profound increase in viral replication in elite controllers [16], followed by a recovery of viral control once CD8+ T cell population was restored [3, 90]. CD8+ T cell responses can also differentiate between rebounders and controllers in SIV-infected macaques, with individuals able to control the infection displaying a long-term memory CD8+ T cell response with enhanced suppressive activity [91].

However, despite the importance of CD8+ T cells in HIV control, their mechanism of action remains poorly-understood [92]. Evidence points to a noncytolytic effect of CD8+ T cells: two groups examined the duration of SIV-infected cell lifespan following treatment with nucleotide reverse transcriptase inhibitors (NRTIs), with or without the presence of CD8+ cells, finding no difference in average cell-lifespan [93, 94]. Subsequent studies have both supported [95–98] and challenged this effect pointing to cytotoxic activity [99, 100]. More recently, mathematical modeling of data derived from SIV-infected macaques treated with the integrase inhibitor ralegravir, in the presence or absence of CD8+ cells revealed a dual effect: before viral integration, the half-life on infected cells was shorter for the group with CD8+ cells compared to the group whose CD8+ cells were depleted. Furthermore, viral production rates were higher in the absence of CD8+ cells. These results suggest that CD8+ cells have both a noncytolytic effect in suppressing viral production, as well as a cytolytic effect before viral integration [87]. Recent studies in vitro have also pointed to a noncytolytic effect of CD8+ T cells by preventing viral production rather than viral infectivity, which is consistent with Policicchio et al [87], as well as with our observation for a tendency toward increased infectivity during rebound [101].

An alternative hypothesis that could explain the increased killing rate of infected cells is natural killer (NK) cell activity. NK cells have been shown to have an antiviral effect in HIV infection [77] and may play an important role in the outcome of the infection [102–104]. Although considered as nonspecific, innate effector cells, there is growing evidence that supports adaptive, memory-like features of NK cells [42]. In humans, this subset of long-lived NK-cells with memory-like features has been shown to exhibit increased effector functions upon re-encountering viral antigens or proper activation with pro-inflammatory cytokines [43–48]. The presence of antigen-specific NK cells has also been reported in rhesus macaques infected with SIV/SHIV. These cells exhibit specificity towards gag- and env-viral proteins loaded on autologous DCs [105]. Notably, this response appears to be long-lasting, since animals vaccinated with an adenovirus 26-vectored SIV vaccine 5 years prior still showed a recall of these adaptive traits in NK cells, associated with a high degree of cytotoxicity. Nevertheless, this observation lacks direct evidence of NK cell memory responses in humans [106]. In humans, the temporal efficacy of NK cell activity during HIV infection remains not well-understood. Some studies have shown increased cytolytic activity of NK cells during HIV infection [107]. Even though HIV-1 infection initially causes a substantial decrease in the NK cell population, their function of those NK cells is enhanced during viremic infection, compared to treated and suppressed individuals or negative controls [108]. However, it has also been shown that NK cells upregulate inhibitory receptors during viremic HIV infection, affecting the killing ability of NK cells [109, 110], as well as cause sustained impairment [111]. Considering the importance of innate responses in infant immunity [31, 34, 49], as well as a bias of Th2 responses in infants [112], the ability of NK cells contribute to cytolytic effects during pediatric HIV infection is a valid hypothesis that should be further investigated.

Even though we find that only one of the selected models predict a difference in the mean viral clearance, the distribution of individual values for rebound is wider and includes values consistent with rebound of HIV adult infections [57, 73]. Counterintuitively, this difference is not attributed to neutralizing antibody responses but to innate responses, since the distribution of parameter values does not differ on whether the RMs had developed neutralizing antibodies (Supplementary Fig 3 and 4). The role of humoral responses in viral control in infants is poorly examined [3]. In adults, the frequency of HIV-specific memory B cells in elite controllers is higher compared to individuals on ART [3, 113]. Although this has not been examined in pediatric ECs, B cell dysfunction appears to be a distinctive feature of HIV disease advancement in children [3, 114]. In addition to neutralization, antibodies capable of antibody-dependent cell-mediated cytotoxicity (ADCC) have been identified in pediatric non-progressors [3, 115], as well as adult ECs. However, we do not observe differences in viral clearance rate among RMs based on whether they had developed antibodies capable of ADCC activity (data not shown).

There are aspects of the infant immune system that may be utilized for an HIV cure or prevention: for example, infants develop broadly neutralizing antibodies earlier and with higher breadth and potency compared to adults [116–119], with some bnAbs targeting up to 4 different epitopes [120, 121]. Taken together with our results, it appears that cellular immunity plays a vital role in HIV control [122], as we observe an increase in parameters related to CD8+ T cell responses with the role of antibodies being promising but less clear, as indicated by the increased viral infectivity suggested by a large proportion of our model ensemble. Considering recent advances in the study of NK cells, their effect in HIV infection should be further elucidated. In addition, our results further suggest that approaches that induce both B and T cell responses, could augment cellular and humoral immunity to act synergistically for HIV control or prevention. Experiments that take measurements of all these compartments (antibodies, NK cells, CD8+ T cells) in the same individuals may help shed light to these complex dynamics.

## Acknowledgements

J.M.C. acknowledges the support of the National Science Foundation (grant no. DMS-1714654) and National Institutes of Health (grant nos. R21-AI143443-01A1 and R01-OD011095). E.M. acknowledges the support of the National Science Foundation (grant no. DMS-1714654). G.M.S acknowledges the support of the National Institutes of Health (grant no. R01 AI160607). C.C. acknowledges the support of the National Institutes of Health (grant no. R25AI140495). S.R.P. acknowledges the support of the National Institutes of Health (grant no. P01-AI131276-05).

## Notes

### Competing Interest Statement

Dr Conway has served as a consultant for Excision BioTherapeutics and Merck. Dr. Permar serves a consultant for Moderna, Merck, Pfizer, GSK, Dynavax, and Hoopika on their CMV vaccine program and has led a sponsored program with Moderna and Merck on CMV vaccines.

