## Supplementary material for "Comparative Analysis of Within-Host Dynamics of Acute Infection and Viral Rebound Dynamics in Postnatally SHIV-Infected ART-Treated Infant Rhesus Macaques": B2AcRe Supplementary Figures.docx

Supplementary Information

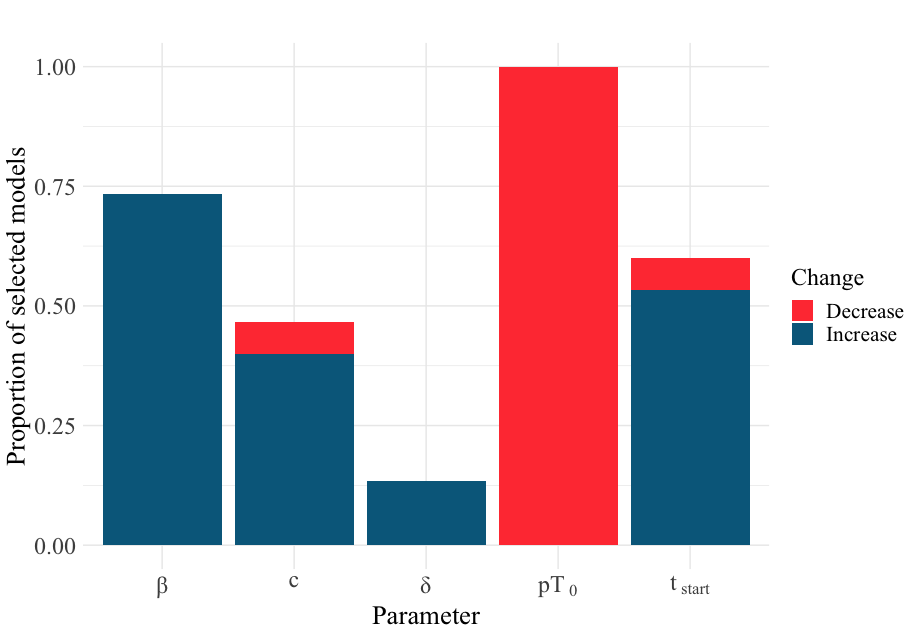

**Supplementary Fig 1:** Bar chart depicting the proportion times of a parameter is predicted to differ between acute infection and viral rebound in all selected models with ΔAIC<8 and *S*<80. A total of 15 models were selected. Viral clearance rate (c) is predicted to be different between acute infection and viral rebound in 7 models; death rate of infected cells (δ) in 2 models; start of exponential viral growth (t_start_) in 9 models; mass-action infectivity (β) in 11 models and the product of viral production rate (p) and start of initial number of target cells (T_0_) in all 15 models. Red slices signify a decrease in the value of the parameter for rebound, whereas blue-toned slices depict an increase in the rebound value of the parameter. Allowing for an increased *S* threshold introduces variability in the predicted association between the stage of the infection and the mean value of the viral clearance rate and the onset of viral exponential growth.

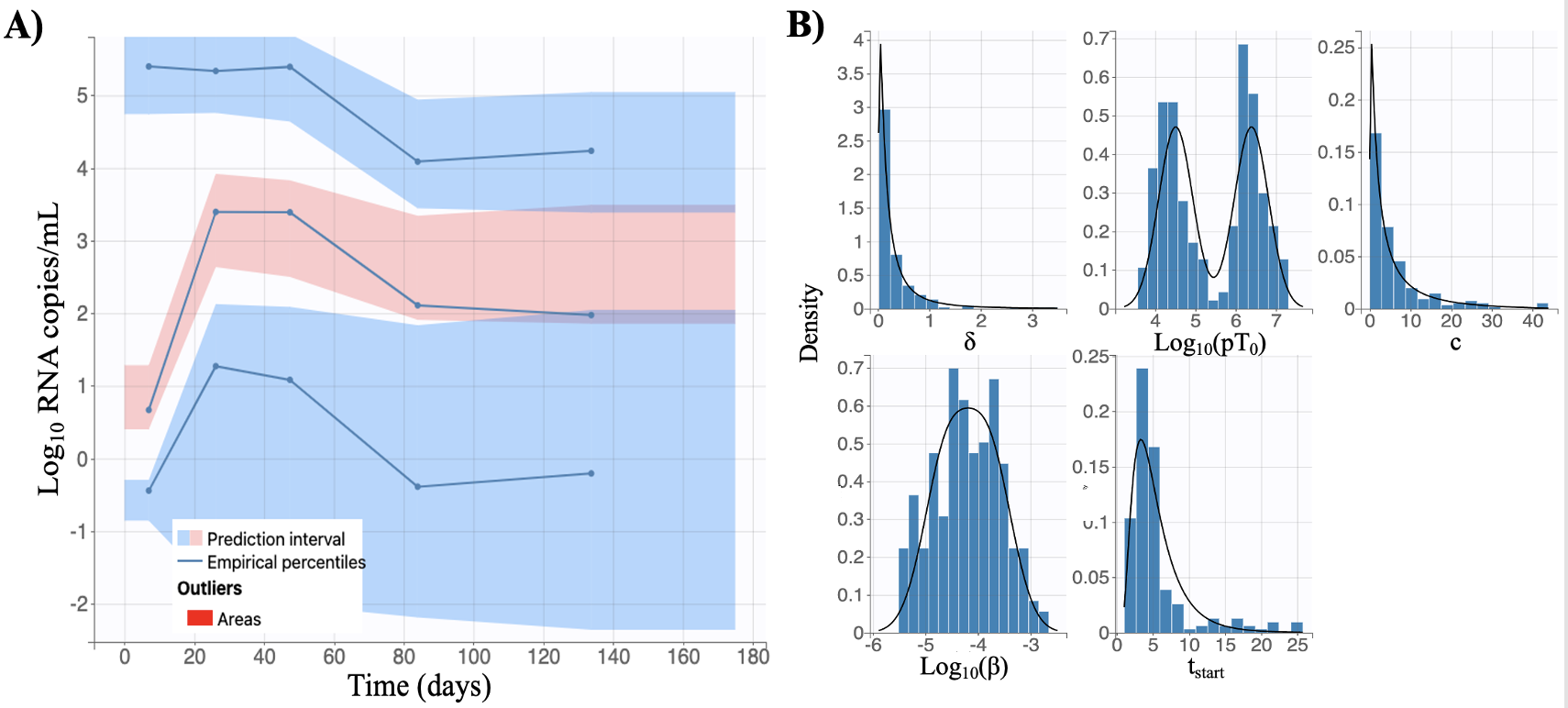

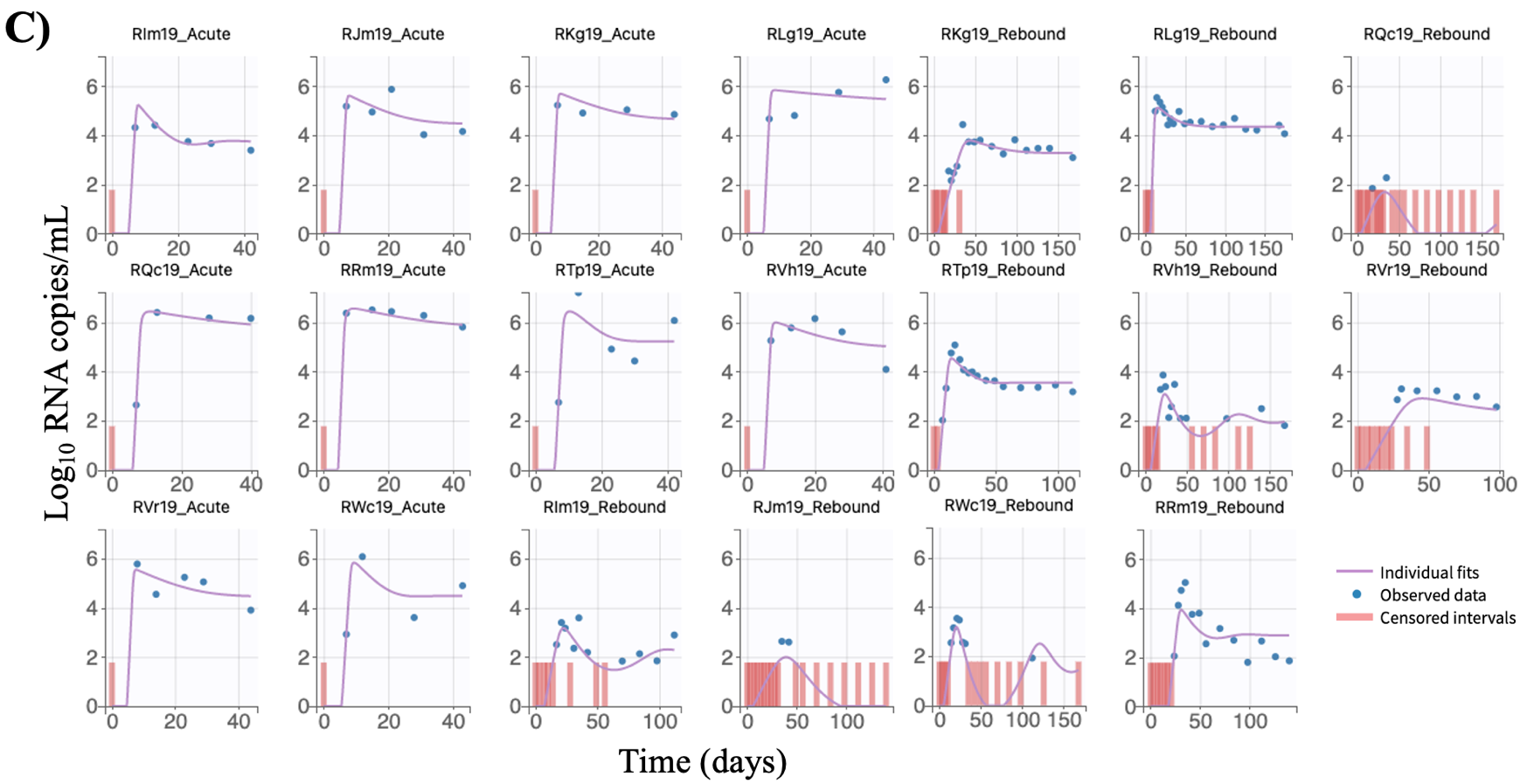

**Supplementary Fig 2:** Example of one of the selected models selected. In this model we fix the median viral clearance rate to c=3 day^-1^for acute infection, and we assume that the product of viral production rate and initial number of target cells (pT_0_) as well as the mass action infectivity (β) are difference between acute and rebound and add a correlation between c and β. A) Visual predictive check estimated from the model. B) Empirical distribution of estimated individual parameter values (blue bars) along with theoretical distribution (black line). C) Viral load measurements for each RM (points) along with the fitted curve (line). Measurements below the limit of detection are depicted in red bars.

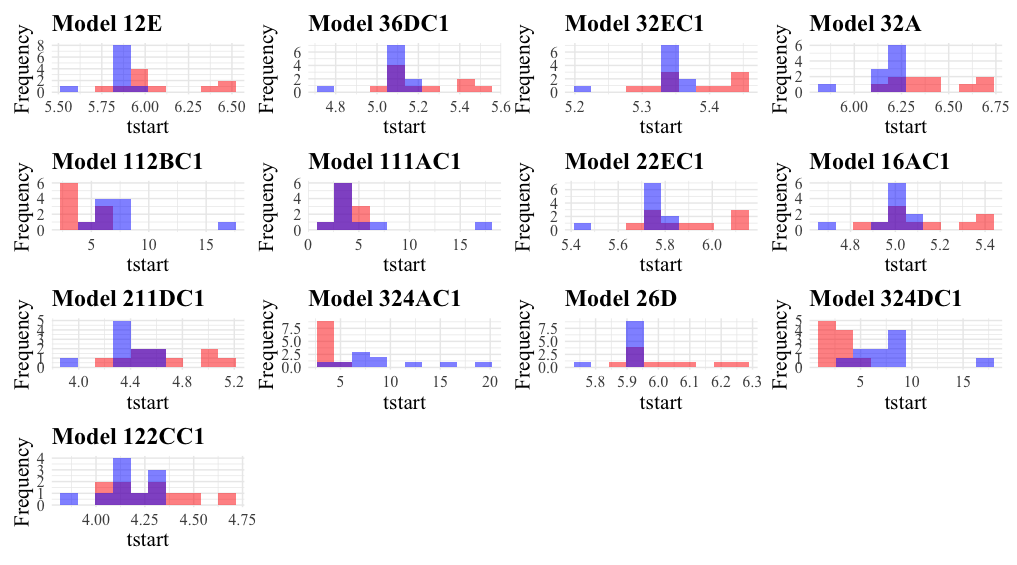

**Supplementary Fig 3:** Distribution of the onset of exponential viral growth (t_start_) for acute infection (red) and viral rebound (blue) for all selected models.

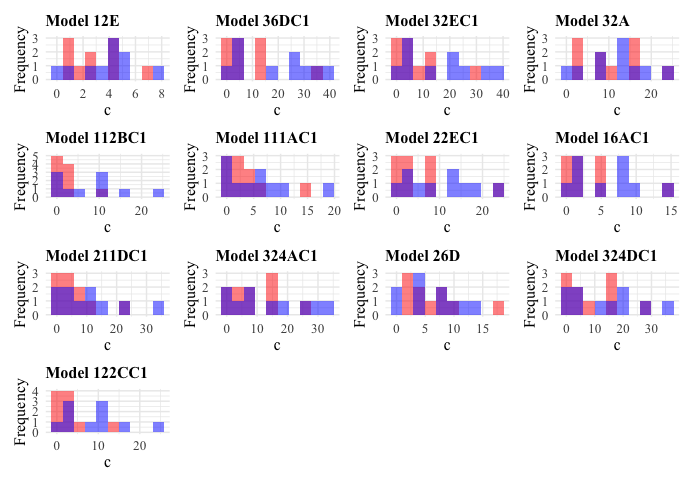

**Supplementary Fig 4:** Distribution of the estimated viral clearance rate (c) for acute infection (red) and viral rebound (blue) for all selected models.

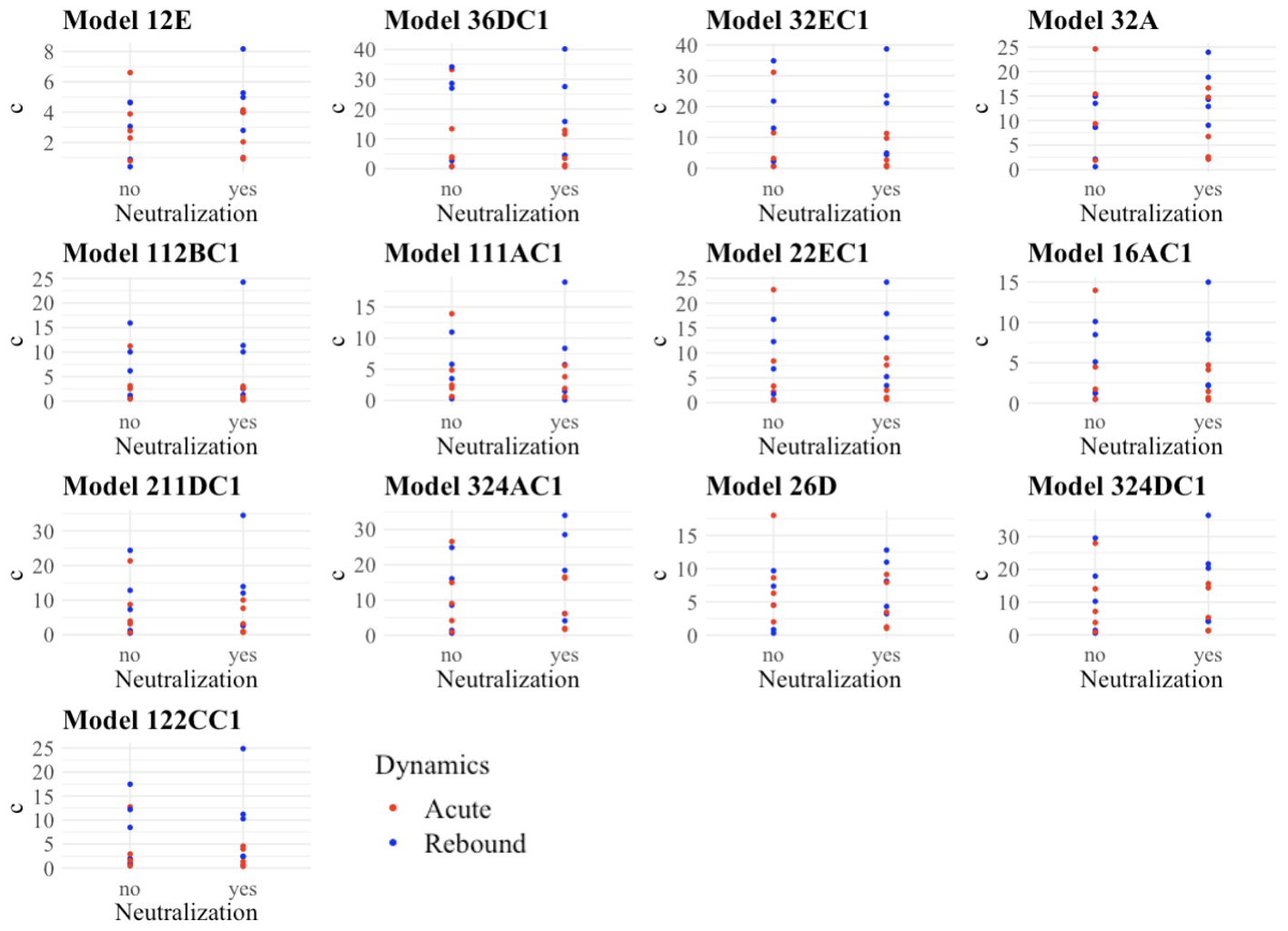

**Supplementary Fig 5:** Scatterplot of the estimated viral clearance rate per day (c) for acute infection (red) and viral rebound (blue) for all selected models based on whether the RM developed antibody neutralizing responses.

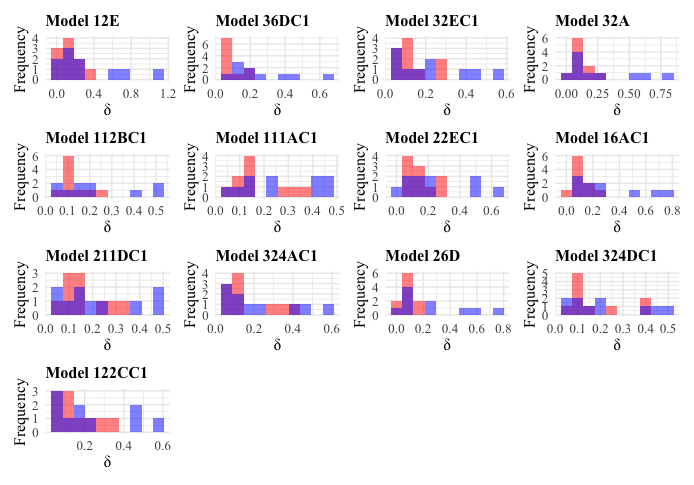

**Supplementary Fig 6:** Distribution of the estimated infected cell death rate (δ) for acute infection (red) and viral rebound (blue) for all selected models.

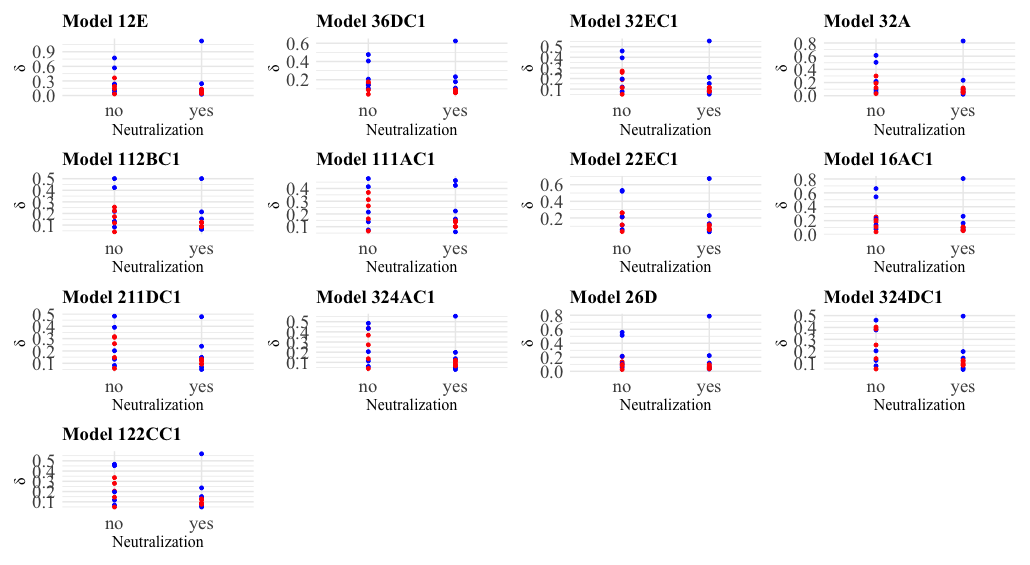

**Supplementary Fig 7:** Scatterplot of the estimated death rate of infected cells per day (δ) for acute infection (red) and viral rebound (blue) for all selected models based on whether the RM developed antibody neutralizing responses.
